# Suppression of Docetaxel induced apoptosis in Survivin-depleted cells due to failure of sustained mitotic block

**DOI:** 10.1101/834259

**Authors:** Teng-Long Han, Hang Sha, Zhi-Xin Jiang

## Abstract

The antitumor effect of taxanes have been attributed to their ability to induce mitotic arrest through activation of the spindle assembly checkpoint. Cell death following prolonged mitotic arrest is mediated by the intrinsic apoptosis pathway. Accordingly, factors that influence the robustness of mitotic arrest or disrupt the apoptotic machinery might confer drug resistance. Survivin is an inhibitor of apoptosis protein. Its overexpression has been associated with resistance to multiple anticancer agents including taxanes, and its targeting led to drug sensitization. On the other hand, Survivin is a key regulator of mitosis, which is shown to be required for stable activation of the spindle assembly checkpoint. Since the sensitivity of taxanes depends on a functional spindle checkpoint, inhibition of Survivin may lead to drug resistance, which is the opposite effect of its anti-apoptotic function. Here we show that Survivin-depleted cells escape the mitotic block following Docetaxel treatment, thereby evading drug induced apoptosis. Moreover, Survivin depletion increases the level of mitotic catastrophe and cellular senescence induced by Docetaxel and enhanced its efficacy against clonogenic survival of tumor cells. Our finding suggests that inhibition of Survivin promotes non-apoptotic mechanisms following Docetaxel treatment rather than increases the sensitivity of apoptosis.

## Introduction

Docetaxel and paclitaxel are taxanes widely used in clinic for the treatment of ovarian, breast, lung cancer and other solid tumors ^1^. They share a common mechanism of action by promoting and stabilizing microtubule assembly, which disrupt microtubule dynamics and chronically activate the spindle assembly checkpoint (SAC), thereby inducing a prolonged mitotic arrest that ultimately leads to cell death ^2^. Although mechanism unrelated to mitosis has been reported and paclitaxel also enacts cell death in interphase^3^, pro-apoptotic signals are frequently found to accumulate during a prolonged mitotic arrest ^4^. Moreover, anti-apoptotic proteins often decrease during mitotic arrest or lose their pro-survival functions ^5^. Therefore taxane induced mitotic arrest is strongly associated with apoptosis. Premature mitotic exit, also known as mitotic slippage, usually due to a defective SAC is considered a process through which tumor cells evade killing by taxanes ^6^. In addition, overexpression of anti-apoptotic proteins also constitutes an important mechanism underlying drug resistance and inhibitors of pro-survival Bcl-2 family members have been shown to enhance taxane induced cell killing ^7^.

Survivin, the smallest member of the inhibitor of apoptosis (IAP) family, is thought to function in both cell cycle regulation and apoptosis inhibition ^8^. Unlike other IAPs, Survivin shows a clear cell-cycle dependent expression at mitosis and degrades by the ubiquitin-proteasome pathway in the G1 phase of the cell cycle ^9^. Together with Aurora-B kinase, INCENP, and Borealin/Dasra, Survivin forms as component of the chromosome passenger complex (CPC) that is essential for proper chromosome segregation and cytokinesis ^10^. Depletion of Survivin invariably results in aberrant mitotic progression ^11^ and its mitotic phosphorylation by p34cdc2-cyclin B1 has been associated with increased stability ^12^.

The role of Survivin in apoptosis inhibition has been controversial. Although Survivin is unable to directly bind to and inhibit caspase activity like other IAPs ^13^, over-expression of Survivin has been associated with the inhibition of cell death induced by multiple anticancer agents ^14,15^. Furthermore, Survivin expression is undetectable in normal adult tissues but selectively expressed in transformed cells and in most human cancers, which make Survivin a promising therapeutic target for the treatment of cancer ^9^. Survivin targeting with antisense nucleotides, ribozymes, or expression of dominant-negative mutants resulted in spontaneous caspase-dependent cell death ^16^ and tumor sensitization to chemotherapeutic agents, such paclitaxel ^17^, Docetaxel ^18^, cisplatin ^19^ and etoposide ^20^.

On the other hand, as an essential mitotic regulator, Survivin has been shown to be required for stable SAC activation ^21,22^. Therefore, Survivin-depleted cells might fail to undergo sustained mitotic arrest following taxane treatment and develop drug resistance, which is contrary to the function of Survivin as a chemo-sensitizer reported in the literature. In this study, we sought to clarify the role of Survivin in cellular response to Docetaxel, using different cell lines, incorporating lentiviral mediated Survivin overexpression and siRNA-mediated Survivin depletion.

## Results

### Cell death following Docetaxel treatment is closely linked with mitotic arrest

By promoting and stabilizing microtubule assembly, taxane treatment activates the SAC and lead to mitotic arrest of tumor cells. However, the choice of cell line, as well as drug concentration, could dramatically affect the treatment response ^4^. Cervical carcinoma HeLa cells or breast cancer MDA-MB-231 cells were treated with different concentration of Docetaxel. While higher concentration of the drug lead to greater cell death in HeLa cells, MDA-MB-231 cells were significantly more resistant to high concentration of Docetaxel (Figure 1A). To explore the mechanisms of resistance, we determined the duration of mitotic block in these two cell lines. HeLa cells exhibited sustained mitotic arrest following treatment of 256nM Docetaxel. In contrast, MDA-MB-231 cells escaped mitotic arrest rapidly (Figure 1B). At 15 hr treatment, both cell lines exhibited typical mitotic morphology with condensed chromatin. While HeLa cells remained in mitosis after 24 hr drug treatment, MDA-MB-231 cells manifested chromosome de-condensation and multi-nucleation, indicative of mitotic exit (Figure 1C).

**Figure 1.**
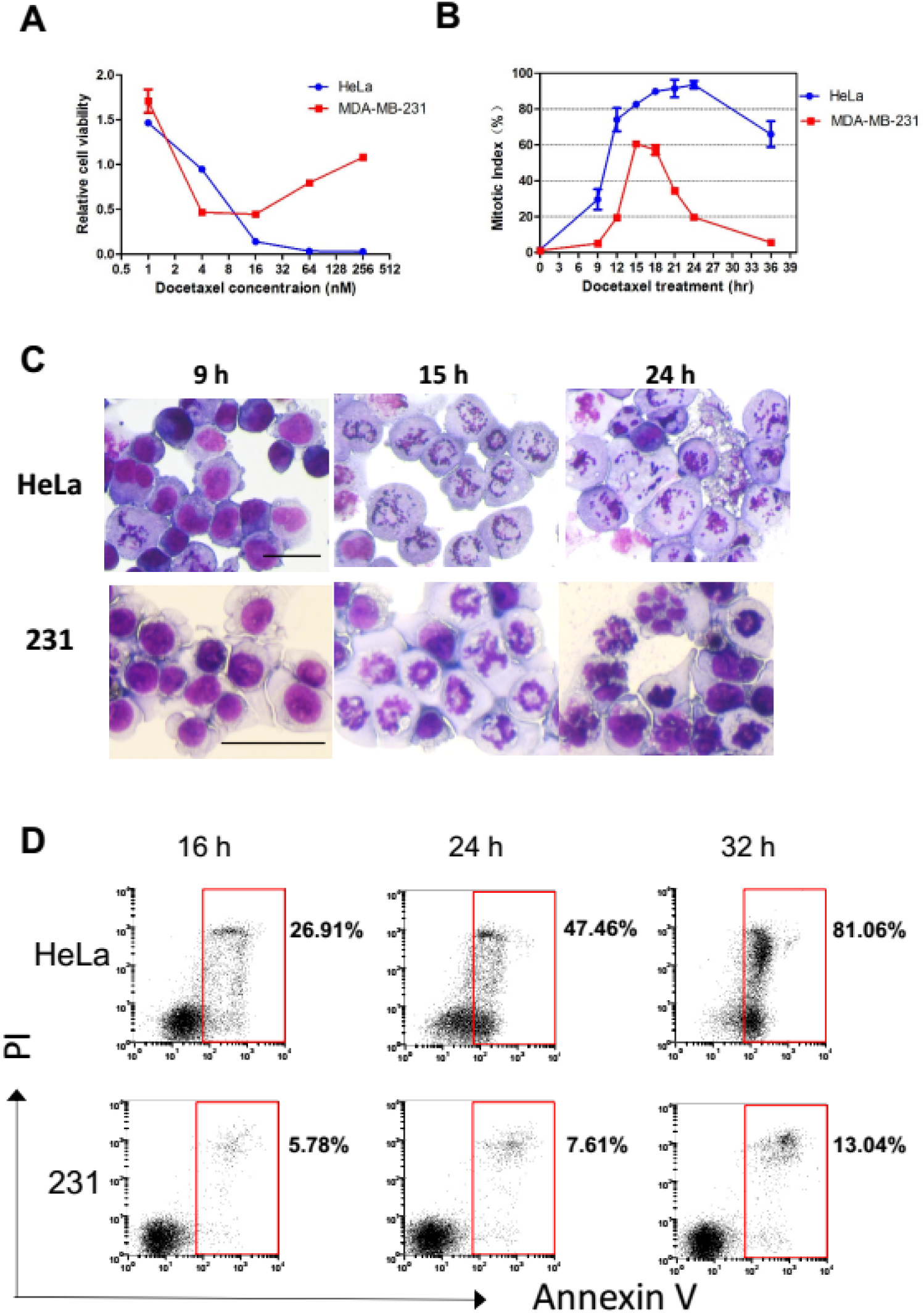
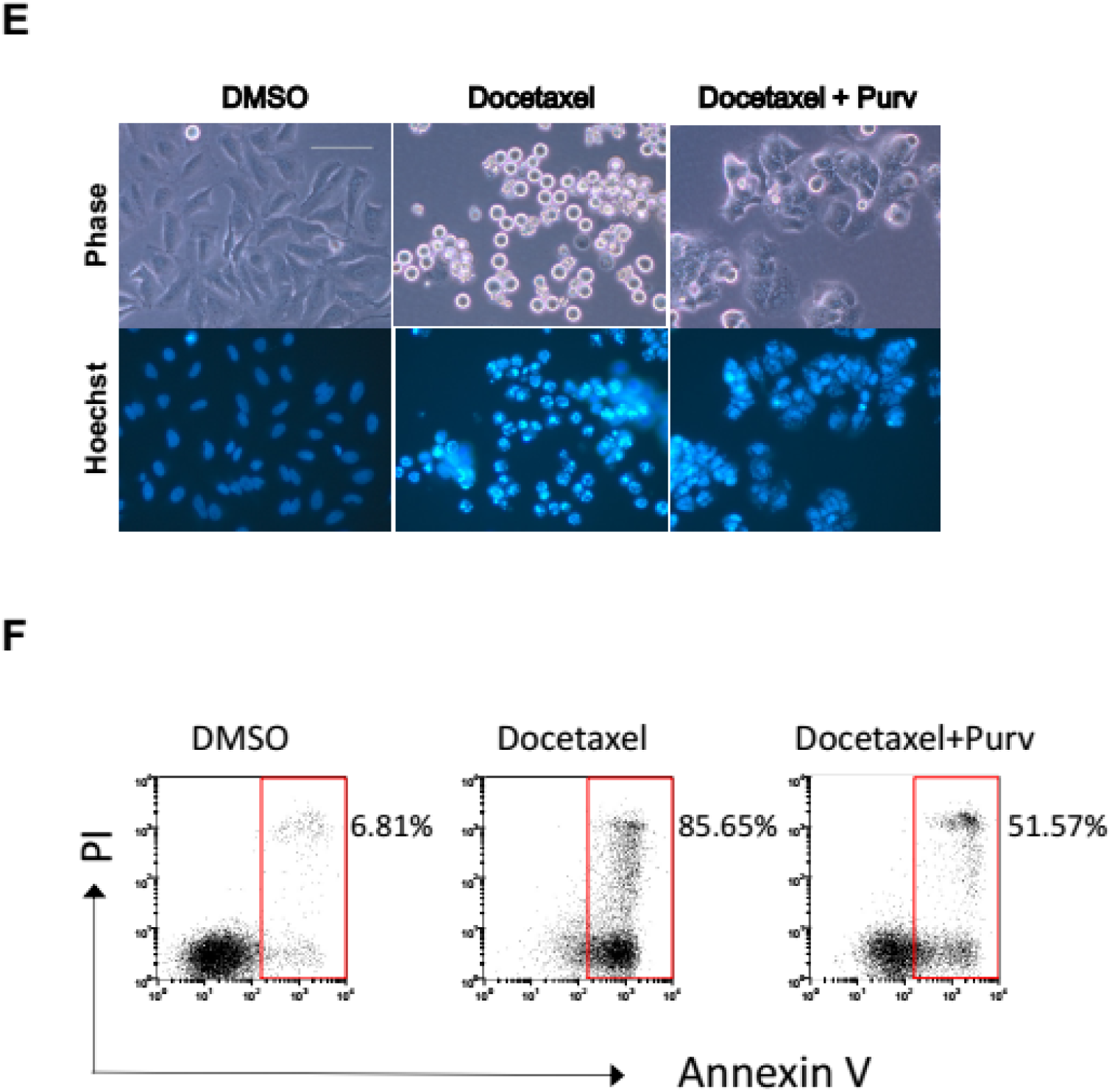
Cell death following Docetaxel treatment is closely linked with mitotic arrest. (A) Cell viability assay. HeLa and MDA-MB-231 Cells were treated with indicated increasing concentrations of Docetaxel, cell viability was determined 3 days following treatment. Values represent the means ± S.D. (n = 3 wells). (B) Mitotic index. Cells were synchronized at the G1/S boundary by double thymidine block. Upon release from the block (0 hr), the cells were treated with Docetaxel (256 nM) and collected at indicated time intervals. Cells were then subjected to Giemsa staining. Mitotic indexes were determined by counting the percentage of cells with condensed chromatin. Values represent the means ± S.D. (n = 3 fields). (C) Mitotic arrest versus mitotic exit. Cells were treated as described in (B). Representative fields of HeLa and MDA-MB-231 cells at the indicated time points are shown. The scale bars represent 50μm. (D) Synchronized cells subjected Docetaxel (256 nM) treatment were collected at indicated time points and stained for Annexin V/PI. Annexin V–positive (apoptotic) cells were analyzed by FACS. (E and F) Synchronized HeLa cells were exposed to Docetaxel (64nM) for 16 hr and then Purv (10 μM) was added for promotion of mitotic slippage. After 30 hr Docetaxel treatment with or without Purv, the cells were stained with Hoechst 33342 and photographed using phase and Hoechst fluorescence (E) or analyzed for induction of apoptosis (F).

The accumulation of pro-apoptotic signaling during prolonged mitotic arrest has been described by many studies ^2,23^. We subsequently determined the level of apoptosis at different time points starting from 16 hr treatment when most HeLa cells became blocked in mitosis. A marked increase in apoptotic cells (from 26% to 81%) during 16 hr was observed in HeLa cells that arrested in mitosis in response to Docetaxel (Figure 1D). In contrast, we only observed a slight increase in cell death (from 5% to 13%) during the same period of time in MDA-MB-231 cells which undergone mitotic exit.

To confirm the association between mitotic arrest and apoptosis, HeLa cells were forced to exit mitosis following 16 hr exposure to docetaxel by addition of purvalanol A (Purv) which inhibits the mitotic kinase Cdk1 thus facilitates premature mitotic exit despite the activation of SAC. Purv treatment yielded large cells with multiple micronuclei and de-condensed chromatin, suggesting cells have aberrantly entered the subsequent interphase (Figure 1E). By contrast, cells treated with Docetaxel alone exhibit spherical morphology and condensed chromatin suggesting mitotic blockage. Notably, the level of apoptosis induced by 30 hr Docetaxel treatment was markedly reduced (from 85% to 51%) by addition of Purv (Figure 1F). Together, these observations suggest that apoptosis induced by Docetaxel is closely associated with mitotic arrest.

### Over-expression of wild type or T34 mutant form of Survivin does not affect sensitivity to Docetaxel or cisplatin

Expression of anti-apoptotic proteins was frequently found to be involved in the chemo-resistant phenotypes of tumors cells ^24^. Moreover, the anti-apoptotic function of Bcl-2 or Bcl-xL were reported to be abrogated through mitotic phosphorylation by p34cdc2-cyclin B1, thus coupling mitotic arrest to apoptosis ^25^. However, still others report that phosphorylation is required for full and potent anti-apoptotic function of Bcl-2 ^26^. On the other hand, the anti-apoptotic function of Survivin has been shown to depend on mitotic phosphorylation to maintain its stability ^12^.

To examine the effect of mitotic phosphorylation on anti-apoptotic functions, we first compared endogenous expression (Figure 2A) and phosphorylation (Figure 2B) of apoptosis regulators in response to Docetaxel treatment. In accordance with previous observations ^12^, we found the expression of Survivin was closely associated with mitotic phase of the cell cycle. A gradual increase in the level of Survivin was observed in HeLa cells which exhibited sustained mitotic block following Docetaxel treatment, though the concomitant apoptosis suggests it was not cyto-protective (Figure 1D). A similar increase was observed in MDA-MB-231 cell following 0-16 hr Docetaxel treatment. However, the level of Survivin decreased in cells subjected to 24 hr drug treatment which have undergone mitotic exit. MDA-MB-231 cells did not express Bcl-2 but expressed higher level of Bcl-xL than HeLa cells. Phosphorylation of these proteins was observed exclusively in cells that arrested in mitosis, consistent with the mitotic phosphorylation of these anti-apoptotic proteins.

**Figure 2.**
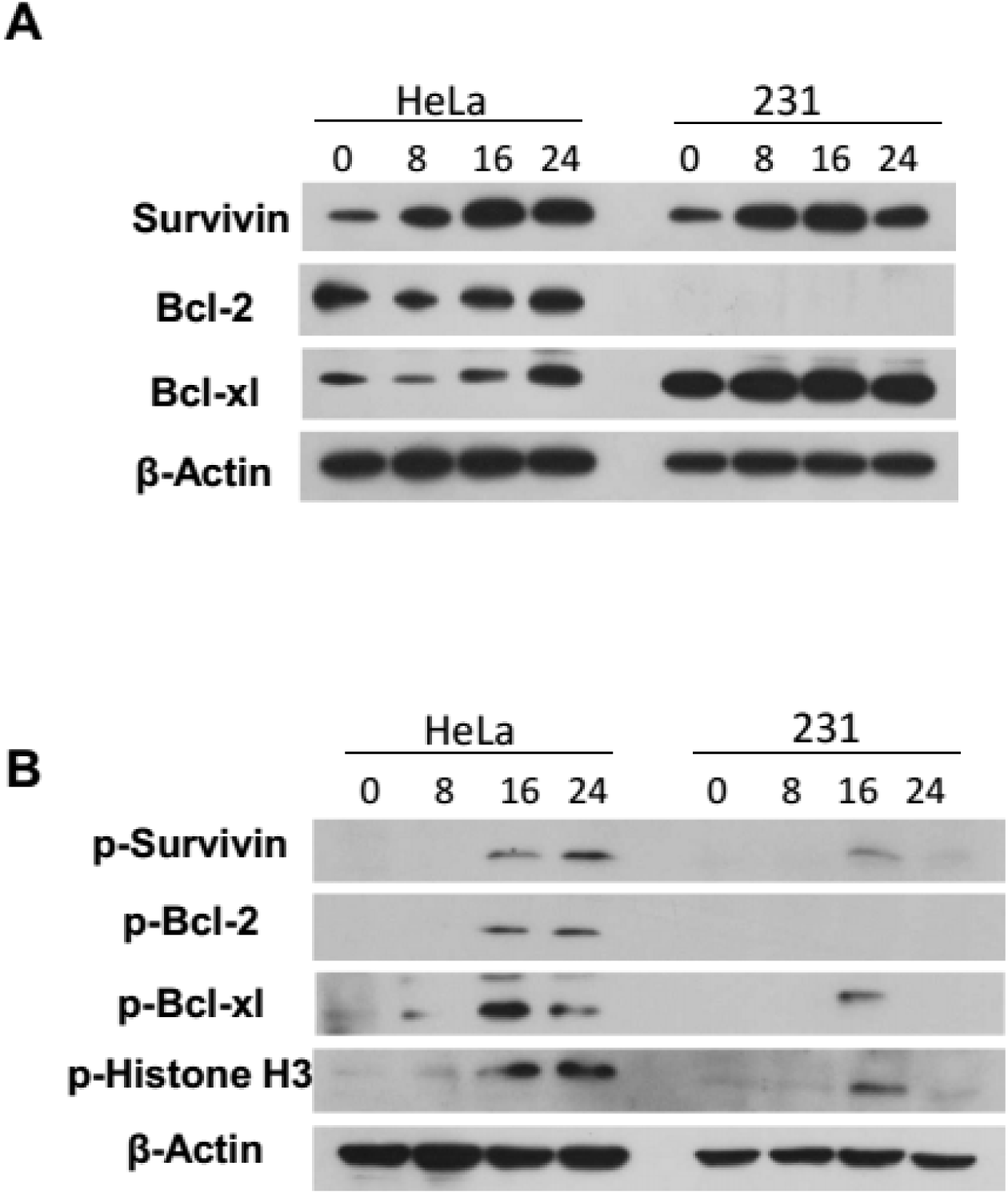
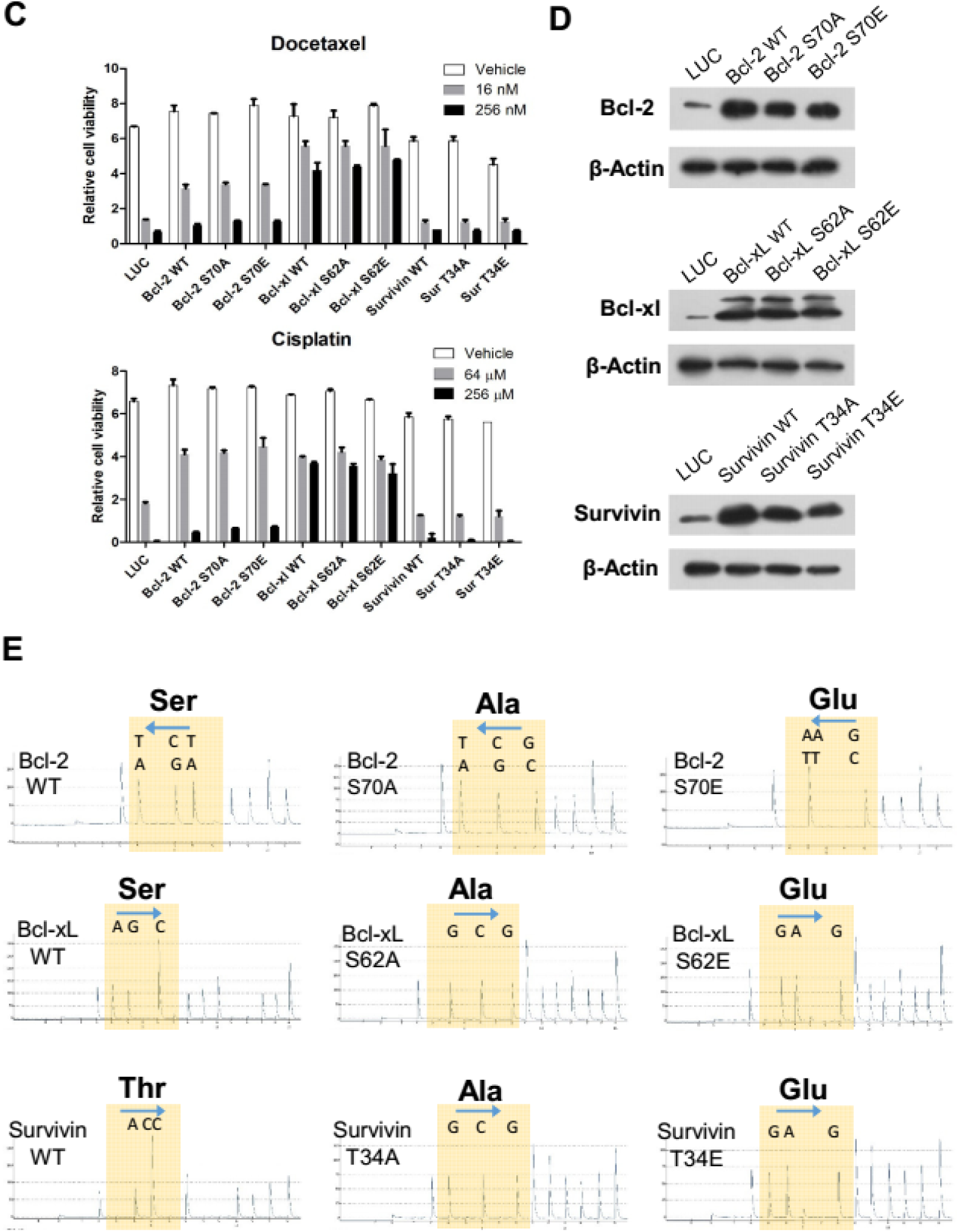
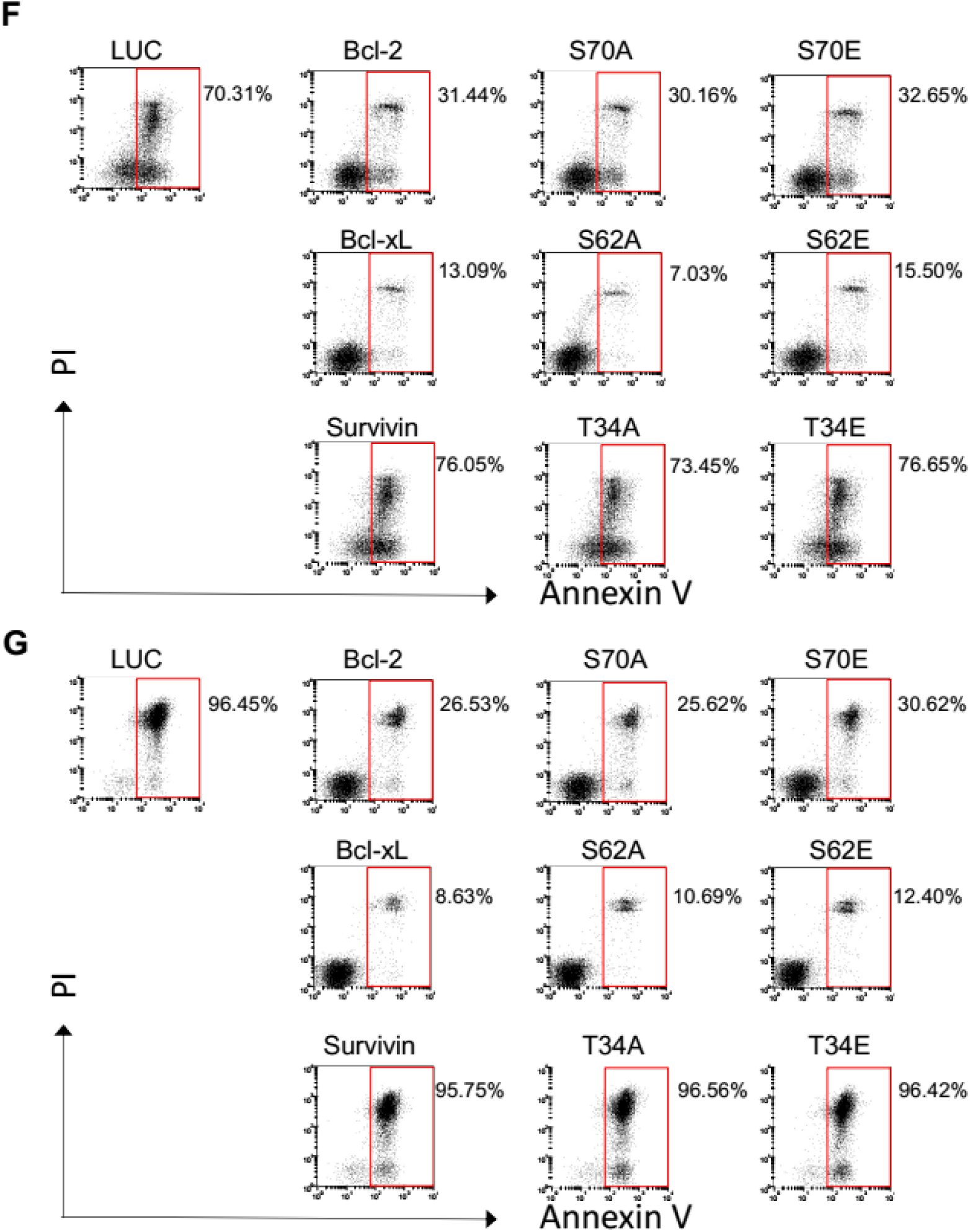
Lentiviral overexpression of Survivin does not confer chemo-resistance. (A and B) Synchronized cells were treated with Docetaxel and harvested at the indicated time intervals. Protein expression (A) or phosphorylation (B) of Survivin, Bcl-2, and Bcl-xL following Docetaxel treatment was determined by western blotting. Phosphorylation of histone H3 was used as a mitotic marker. Original blots can be found in Supplementary Fig. S1. (C) HeLa cells were infected with lentivirus expressing the indicated proteins for 48 h, followed by Docetaxel (upper panel) or cisplatin (lower panel) treatment for another 48 h before cell viability was determined. Values represent the means ± S.D. (n = 2 wells). (D) Expression of indicated protein in stable cell lines was validated by western blotting. Original blots can be found in Supplementary Fig. S2. (E) Expression of mutant protein was validated by cDNA pyrosequencing. (F and G) Stable cell lines expressing the indicated protein were treated with Docetaxel (256 nM) (F) or cisplatin (256 μM) (G), then collected and analyzed for induction of apoptosis.

To express these anti-apoptotic proteins effectively, we took advantage of a lentiviral expression system which circumvent the cytotoxicity mediated by plasmid transfection and shows premium gene transfer efficiency ^27^. HeLa cells were incubated with lentiviral particles expressing Bcl-2, Bcl-xL, Survivin, or phosphomimetic or deficient mutants for 48 hr, followed by treatment with Docetaxel or cisplatin for another 48 hr. Elevated level of cell viability was observed in cells infected with lentivirus expressing Bcl-2 or Bcl-xL, although Bcl-xL seems to be more effective to inhibit the cell death induced by higher concentrations of Docetaxel or cisplatin, in accordance with its more potent anti-apoptotic function ^28^ (Figure 2C). Infection of lentivirus expressing phosphorylation site mutant Bcl-2 or Bcl-xL resulted in similar level of cell viability compared with wild type proteins. Notably, infection of lentivirus expressing of Survivin, whether wild-type, phosphomimetic or deficient mutant, did not show any protective effect against docetaxel or cisplatin treatment.

To further test the role of anti-apoptotic proteins in cellular response to therapy, stable cell lines expressing wild-type or mutant forms of Bcl-2, Bcl-xL, or Survivin were generated through selection with puromycin. Over-expression of anti-apoptotic proteins was validated by western blotting (Figure 2D) and phosphorylation site mutants by pyrosequencing the cDNA of stable cell lines (Figure 2E). When treated with Docetaxel or cisplatin, cells over-expressing Bcl-xL were the most resistant to cell killing induced by either drug (Figure 2F and 2G). The level of apoptosis was also reduced by over-expression of Bcl-2. Notably, their anti-apoptotic activities were not markedly affected by single amino acid mutation at phosphorylation site. This observation does not rule out the possibility that phosphorylation of Bcl-2 or Bcl-xL may regulate their anti-apoptotic function, for the functional differences between wild-type and mutant proteins may be masked by the high level expression of exogenous proteins through lentiviral infection. Moreover, treatment of taxanes was reported to lead to multisite hyper-phosphorylation of Bcl-2 family proteins ^29^. Thus, the effect of mono-phosphorylation might be limited. Nevertheless, our data demonstrated that the anti-apoptotic property was retained in Bcl-2 or Bcl-xL protein with single amino acid mutation at phosphorylation site. Of note, over-expression of wild-type Survivin or its phosphomimetic or deficient mutants showed no effect on cellular sensitivity to either docetaxel or cisplatin treatment.

### Varied toxicity of siRNAs targeting Survivin

Depletion of Survivin has been reported to result in spontaneous apoptosis as well as sensitization to chemotherapeutic agents ^16^. Considering the difference in knockdown efficiency and off-target effect between different siRNA sequences, either commercially designed sequences (#1–3) or a sequence from previous literature (#4) ^21^ were used to target Survivin. All of these siRNAs were effective at knockdown the protein level of Survivin (Figure 3A). However, the level of apoptosis induced by individual siRNA varied widely (Figure 3B). While transfection of siRNA #2 or #3 resulted in substantial apoptosis, the level of apoptosis induced by siRNA #1 or #4 was only slightly more than that of control siRNA.

**Figure 3.**
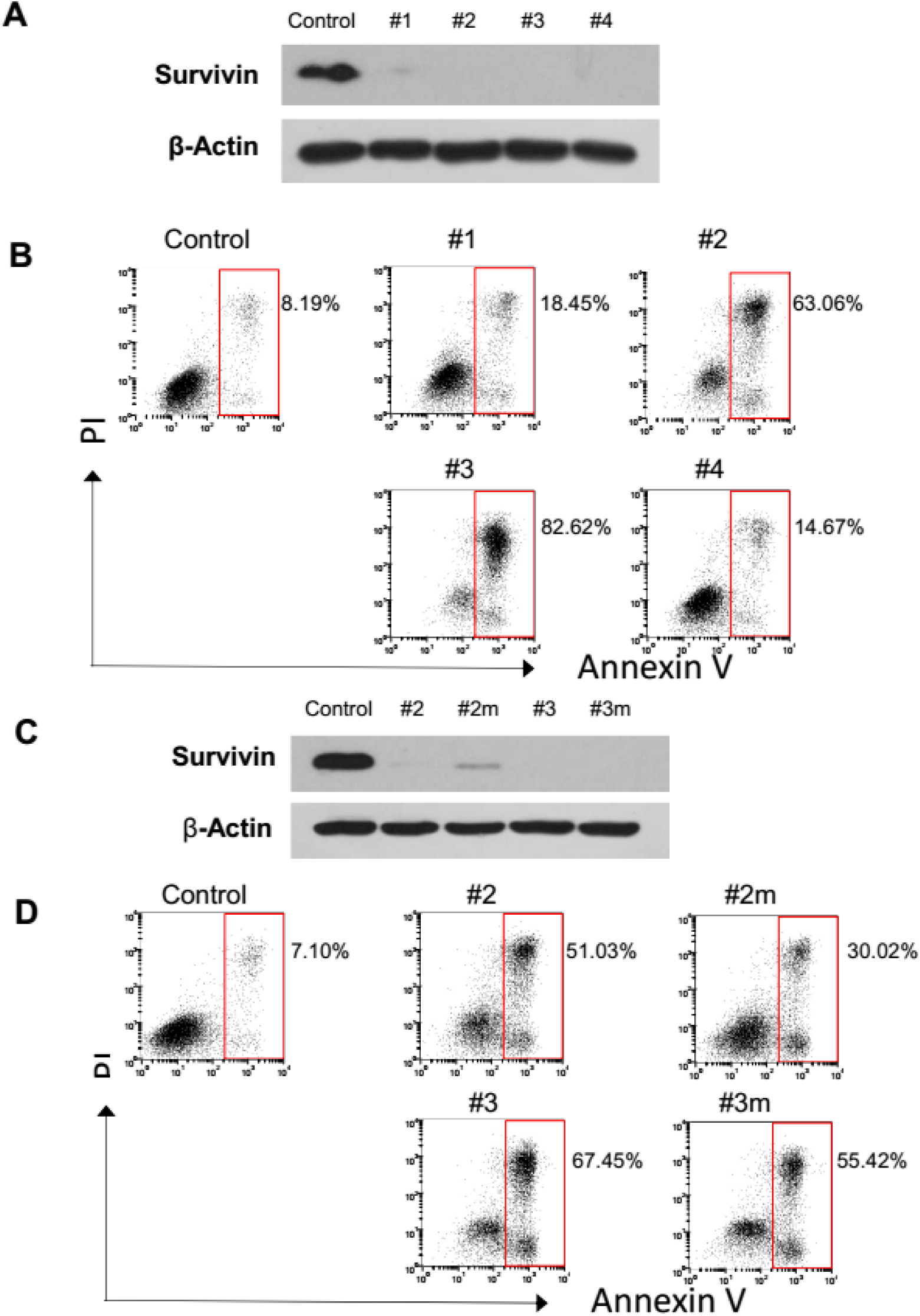
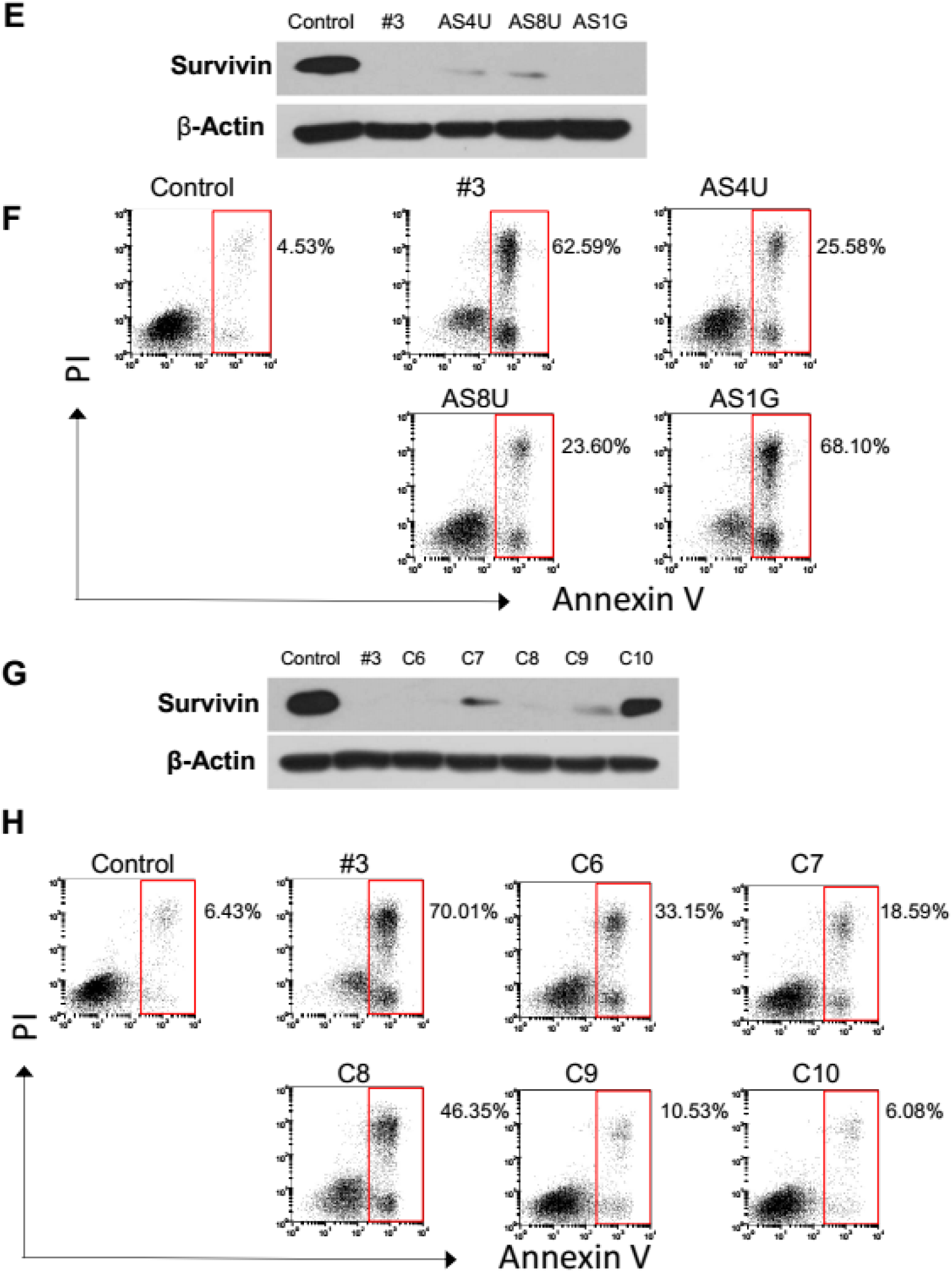
Varied toxicity of siRNAs targeting Survivin. (A) Depletion of Survivin by siRNA transfection. HeLa cells were incubated with indicated siRNA for 48 hr, then harvested and analyzed for Survivin expression by western blotting. Original blots can be found in Supplementary Fig. S3. (B) Different toxicity of siRNAs targeting Survivin. HeLa cells were incubated with indicated the siRNAs for 60 hr, then collected and analyzed for induction of apoptosis. (C) Depletion of Survivin by chemically modified siRNAs. HeLa cells were transfected with original or 2’-O-methyl modified siRNA. Survivin expression was determined by western blotting. Original blots can be found in Supplementary Fig. S3. (D) Diminished toxicity of chemically modified siRNAs. Cells were treated as described in (B). (E) Depletion of Survivin by siRNAs with modification in the antisense strand. Cells were transfected with original siRNA or siRNA with 2’-O-methyl modification in the antisense strand. Survivin expression was determined by western blotting. Original blots can be found in Supplementary Fig. S3. (F) Diminished toxicity of Survivin siRNA by modification in the antisense strand. Cells were treated as described in (B). (G) Depletion of Survivin by siRNAs with single base substitution. Cells were transfected with original siRNA or siRNA with single base substitution. Survivin expression was determined by western blotting. Original blots can be found in Supplementary Fig. S3. (H) Diminished toxicity of Survivin siRNA by single base substitution. Cells were treated as described in (B).

Off-target activity of siRNAs has been shown to result in a toxic phenotype, which is apoptotic in nature ^30^ and could be reduced by position-specific chemical modifications ^31^. Accordingly, we found the toxicity of siRNA #2 or #3 was diminished following 2’-O-methyl modification (#2m or #3m) while the gene silencing was not significantly affected (Figure 3C and 3D). A more effective design is to modify only uridine residues in the antisense strand of siRNA ^32^. We found apoptosis induced by siRNA#3 was markedly reduced through 2’-O-methyl modification of either 8 uridine residues of antisense strand (AS8U) or the 4 uridine residues within seed region of antisense strand (AS4U). Whereas, Modification of guanosine residues (AS1G) showed no effect. Survivin silencing was not markedly affected by siRNA modification (Figure 3E and 3F). Similar results were obtained in MDA-MB-231 cells (Supplementary Figure S4).

To further verify the existence of nonspecific toxic effect, mismatched siRNA C6-C10 were generated by changing bases 6–10 of siRNA #3 to their complement base, respectively. Survivin silencing was not markedly affected by siRNA mismatches except C10 (Figure 3G). Of note, apoptosis induced by siRNA#3 was reduced to varying degrees following base substitution (Figure 3H). Taken together, these observations suggested that nonspecific toxic effect may contributed to the apoptosis following transfection of siRNA against Survivin. The siRNA sequence #4 which exhibit minimal toxicity was used in subsequent experiments.

### Depletion of Survivin diminishes Docetaxel induced apoptosis in cells that exhibit stable mitotic block

Inhibition of anti-apoptotic proteins has been shown to enhance cell killing by chemotherapeutic agents including taxanes ^7^. To determine whether depletion of Survivin results in elevated chemo-sensitivity, MDA-MB-231 cells were transfected with of siRNA against Bcl-xL or Survivin and subjected to Docetaxel or cisplatin treatment. Protein depletion was validated by western blotting (Figure 4A). Whereas depletion of Bcl-xL markedly enhanced the cell death induced by Docetaxel or cisplatin, comparable level of apoptosis was detected between cells transfected with Survivin and control siRNA (Figure 4B). Similarly, depletion of Bcl-xL sensitized HeLa cells to Docetaxel treatment, consistent to its anti-apoptotic function. In contrast, depletion of Survivin exhibited a cellular protective effect against Docetaxel treatment in HeLa cells (Figure 4C). The decreased level of apoptosis in Survivin-depleted cells was more pronounced in response to16nM Docetaxel (from 61% to 39%) compared with 64nM (from 65% to 53%).

**Figure 4.**
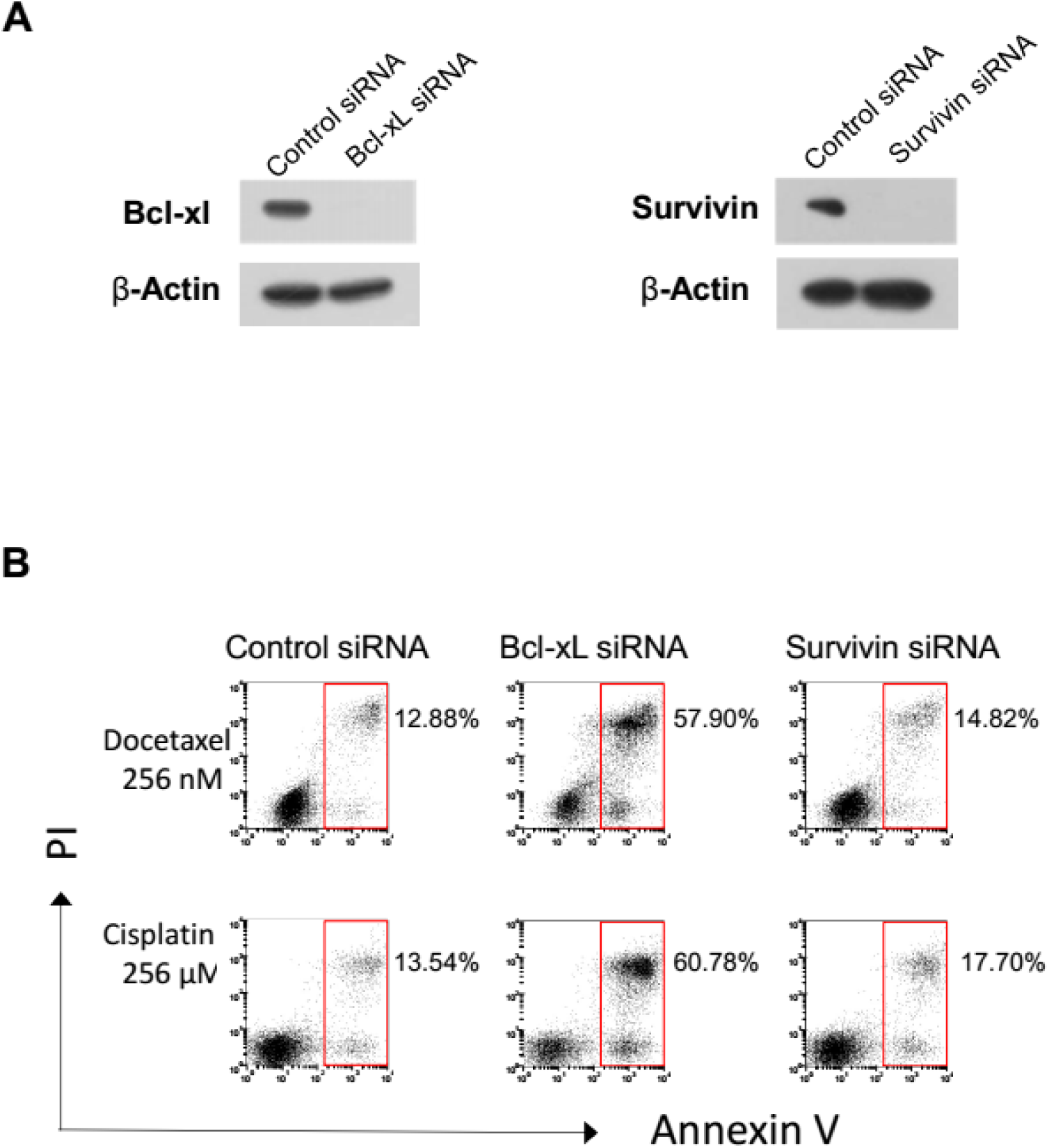
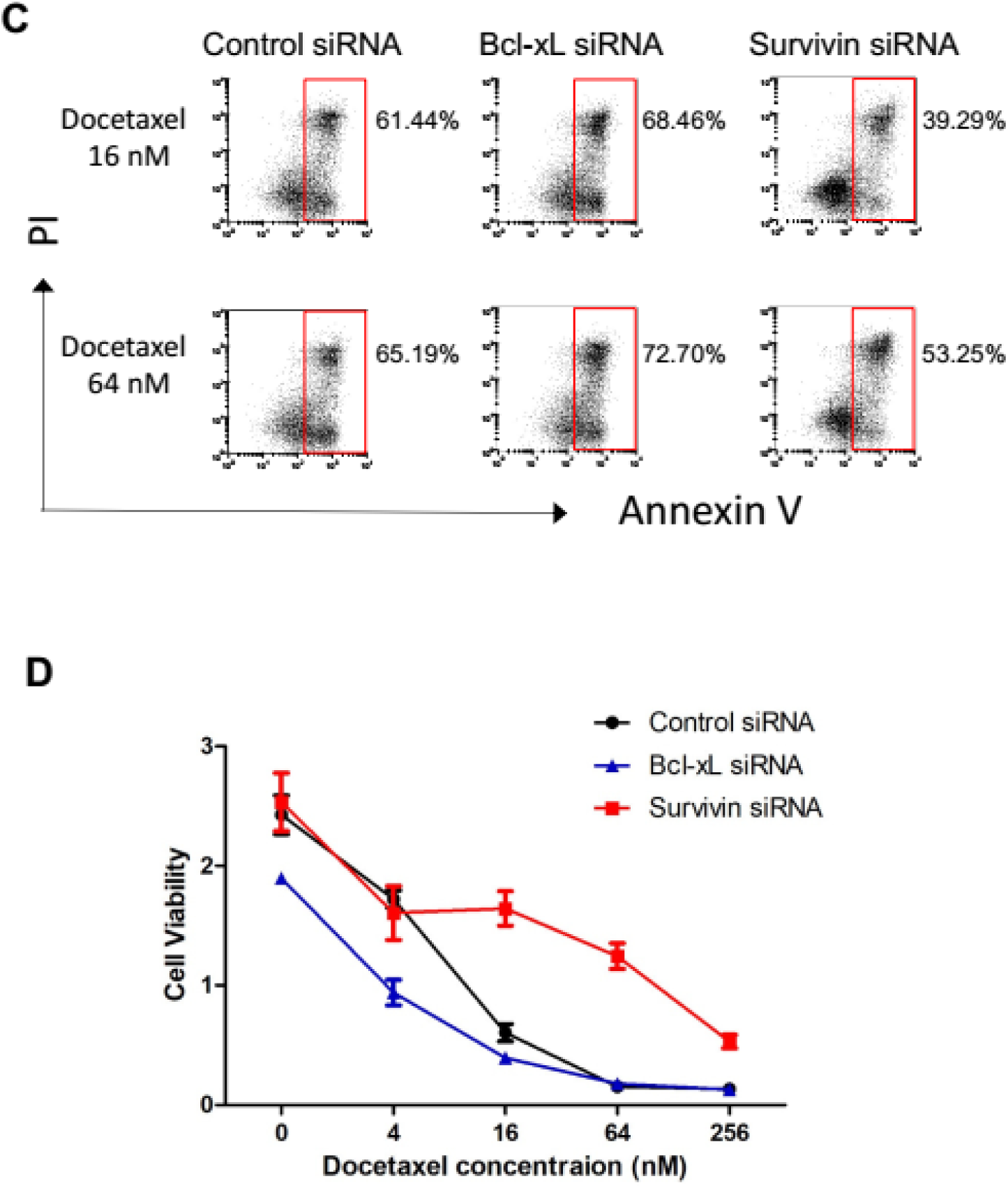
Depletion of Survivin diminishes Docetaxel induced apoptosis in cells that exhibit stable mitotic block. (A) Depletion of Bcl-xl and Survivin by siRNA transfection. MDA-MB-231 cells incubated with the indicated siRNAs for 48 h were harvested and analyzed for Survivin expression by western blotting. Original blots can be found in Supplementary Fig. S5. (B) Sensitization to chemotherapeutic agents by depletion of Bcl-xL but not Survivin. MDA-MB-231 cells were incubated with the indicated siRNAs for 24 hr followed by Docetaxel (upper panel) or cisplatin (lower panel) treatment for 48 h or 24 hr. Cells were collected and analyzed for induction of apoptosis. (C) Depletion of Survivin promoted cell survival against Docetaxel treatment. HeLa cells were incubated with the indicated siRNAs for 36 hr followed by Docetaxel treatment for 24h. Cells were collected and analyzed for induction of apoptosis. (D) HeLa cells were incubated with the indicated siRNAs for 36 hr followed by treatment with increasing concentrations of Docetaxel for 48 hr before cell viability was determined. Values represent the means ± S.D. (n = 3 wells).

To further test the concentration-dependent protective effect of Survivin depletion, we determined viability of Bcl-xL or Survivin depleted cells in response to different concentrations of Docetaxel treatment (Figure 4D). Whereas depletion of Bcl-xl sensitized HeLa cells to different concentration of Docetaxel, depletion of Survivin led to enhanced cell survival. Of note, the protective effect decreased as drug concentration increased. Moreover, depletion of Survivin also led to enhanced cell survival when MCF-7 cells were treated with Docetaxel (Supplementary Figure S6).

### Depletion of Survivin facilitates premature mitotic exit and switches Docetaxel induced apoptosis to senescence

Mitotic slippage due to a defective SAC has been linked with resistance to microtubule toxins. Since Survivin has been reported to be required for sustained activation of the SAC ^21,22^, Survivin-depleted cells tend to undergone mitotic slippage in response to taxanes, and may thereby developing drug resistance. To test this hypothesis, HeLa cells transfected with Survivin or control siRNA were subjected to Docetaxel treatment. Whereas control cells experienced normal mitotic arrest in response to 16 hr treatment of both 16nM and 256nM Docetaxel, mitotic block was only observed when Survivin-depleted cells were subjected to 256nM Docetaxel. However, the cells treated with 16nM Docetaxel showed enlarged cell size, de-condensed chromosome and increased level of multi-nucleation, indicative of mitotic slippage (Figure 5A). Similar observations have been made by a previous study ^33^. These findings suggest that Survivin-depleted cells were capable of escaping the mitotic block induced by low concentration of Docetaxel, thereby avoiding drug induced apoptosis. These data may explain why the protective effect of Survivin depletion is only observed in cells exhibit stable mitotic arrest and is more pronounced in response to low concentration of Docetaxel.

**Figure 5.**
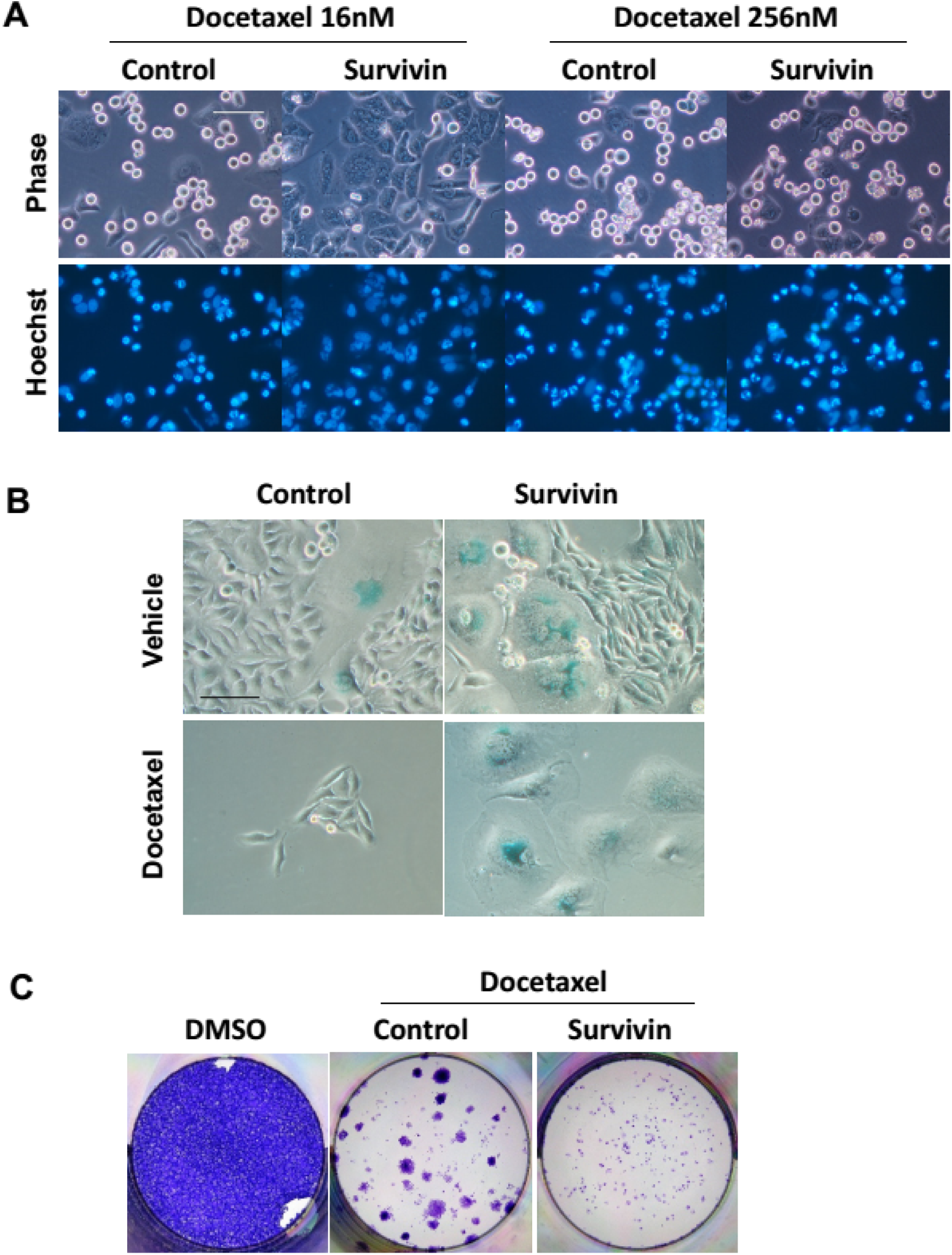
Depletion of Survivin facilitates premature mitotic exit and switches Docetaxel induced apoptosis to senescence. (A) Survivin-depleted cells undergone mitotic slippage when treated with 16nM Docetaxel, but exhibit normal mitotic arrest at 256nM Docetaxel. HeLa cells were incubated with the indicated siRNAs for 36 hr and synchronized by thymidine block. Following a 16 hr exposure to Docetaxel, cells were stained with Hoechst 33342 and photographed using phase and Hoechst fluorescence. The scale bars represent 100 μm. (B) Survivin-depleted cells showed elevated level of mitotic catastrophe and cellular senescence. HeLa cells were incubated with the indicated siRNAs for 48 hr followed by Docetaxel treatment for 36h, and then cultured in drug-free medium. Cellular senescence was determined by SA-β-Gal staining. The scale bars represent 100 μm. (C) Colony outgrowth assay. HeLa cells were incubated with siRNA against Survivin or control siRNA for 48 hr followed by Docetaxel treatment for 36h, and then cultured in drug-free medium for 2 weeks. Cell colonies was photographed following fixation with methanol and staining with crystal violet.

The accumulation of multinucleated giant cells following Survivin depletion conformed to the morphological features of mitotic catastrophe, which has been delineated as a separate mode of cell death. However, accumulating evidences suggest that mitotic catastrophe represents a process preceding apoptosis, necrosis or senescence ^34^. Survivin depletion has been reported to result in mitotic catastrophe ^11^. Here we show that the level of mitotic catastrophe was further increased in response to Docetaxel (Figure 5B). Although Survivin-depleted cells were able to escape the mitotic block induced by Docetaxel, we found proliferative capacity was lost in these cells, which entered a permanent growth arrest state and exhibit morphology features of cellular senescence, as manifested by β-galactosidase (SA-β-gal) activation. In contrast, senescent cells could hardly be observed in control group which mainly undergone apoptotic death in response to Docetaxel. Therefore, Survivin depletion switches Docetaxel induced apoptosis to senescence.

To examine the long–term effect of Survivin depletion on cellular response to Docetaxel, clonogenic assay was carried out which measure the net result of multiple forms of death ^35^. We found the combination of Survivin depletion and Docetaxel showed an elevated efficacy against clonogenic survival of tumor cells compared with Docetaxel treatment alone (Figure 5C), suggesting that inhibition of Survivin might enhance the anti-tumor activity of Docetaxel through promoting non-apoptotic mechanisms such as cellular senescence.

## Discussion

Despite the prevalent use in clinical practice, the mechanism by which taxanes kills tumor cells remains to be clarified. Taxanes work by perturbing spindle assembly, which activates the SAC and lead to a prolonged mitotic arrest. Although the molecular link between mitosis and cell death remains obscure, accumulating evidence suggests that pro-apoptotic signals accumulate during prolonged mitotic arrest, therefore associates mitotic arrest with apoptotic cell death ^2,5^. For instance, DNA damage was found to accumulate during a protracted mitotic arrest which may eventually activates intrinsic apoptotic pathways ^36^. Moreover, anti-apoptotic members of Bcl-2 family, including Bcl-xL, Bcl-2 and Mcl-1, were expected to undergo mitotic phosphorylation by CDK1-cyclin B1. Phosphorylation of Mcl-1 facilitate its poly-ubiquitination and proteasomal degradation ^37^. The anti-apoptotic functions of Bcl-2 and Bcl-xL were reported to be abrogated through mitotic phosphorylation ^25^. Along this line, targeting mitotic exit has been proposed as a powerful therapeutic approach, which results in tumor regression due to apoptotic cell death ^38^.

Although our results demonstrated that anti-apoptotic ability was retained in Bcl-2 or Bcl-xL protein with single amino acid mutation at phosphorylation site, we also observed the marked increase in the level of apoptosis during Docetaxel induced mitotic block. Furthermore, cell death was diminished following treatment of Purv, an inhibitor of CDK1, which promote mitotic slippage. Our observations are in line with the close link between mitotic arrest and apoptotic cell death, suggesting that mitotic slippage could be a mechanism through which tumor cells evade drug-induced apoptosis, therefore, may potentially develop drug resistance.

Overexpression of anti-apoptotic proteins represent another important mechanism underlying drug resistance. Accordingly, we found cell death following Docetaxel or cisplatin treatment was markedly reduced by expression of exogenous Bcl-xL or Bcl-2. Unexpectedly, however, overexpression of Survivin showed no effect on cell survival following treatment. Survivin is a bi-functional protein that acts as both a mitotic regulator and an apoptosis inhibitor ^9^. Its anti-apoptotic function has been a subject of controversy. Over-expression of exogenous Survivin has been associated with chemotherapeutic resistance in multiple studies. We noticed that most observations were based on plasmid transfection or adenoviral infection, which exhibit dose-dependent toxicity to commonly used cancer cell lines ^39,40^. To circumvent unspecific cytotoxicity, we took advantage of a lentiviral expression system which shows premium gene transfer efficiency with minimal cytotoxic effects. While adenovirus infection will finally lead to lysis of the target cells to release viral particles, lentiviruses rely on perturbing cell function as little as possible for their own survival. Owing to the different nature, lentiviral vectors cause little or no disruption of the target cell even at high titer ^27,41^. We found lentiviral overexpression of WT Survivin does not lead to chemo-resistance and overexpression of T34A mutant does not sensitized tumor cells. Consistent with our observations, a study utilizing conditional knockout cells emphasized the role of Survivin in mitosis rather than regulation of apoptosis, and they have made a similar observation that phosphorylation on T34 is dispensable for Survivin function ^33^.

Survivin inhibition was reported to result in spontaneous apoptosis and chemo-sensitization ^16^. However, our observations provide evidence that the nonspecific toxic effect of siRNA transfection are clearly involved in apoptosis following Survivin depletion. Firstly, not all siRNA against Survivin result in spontaneous apoptosis. Significant cell death was only observed following transfection of 2 out of 4 Survivin siRNAs. Moreover, the toxicity of Survivin siRNA was significantly reduced following 2’-O-methyl modification or base substitution, which have been reported to effectively eliminate RNAi mediated off-target effect ^31,42^.

The role of Survivin in cell division is unanimously accepted. It is now clear that Survivin is associated with regulators of cytokinesis, such as Aurora B kinase, INCENP, and Borealin. Together they form the chromosomal passenger complex (CPC) which regulates key mitotic events, including activation and maintenance of the SAC ^10^. Since mitotic arrest in response to taxanes depends on a functional SAC ^43^, depletion of Survivin may facilitate mitotic slippage thereby leading to drug resistance. Others have found that Survivin-depleted cells exhibited a normal mitotic arrest in response to high concentration of taxane treatment. However, they tend to undergo mitotic slippage at lower concentrations, indicative of a defective checkpoint response ^33^. Accordingly, we observed a concentration dependent protective effect of Survivin depletion, which is more pronounced in response to low concentration of Docetaxel. Moreover, the protective effect is only observed in cells exhibit stable mitotic arrest. Cellular sensitivity to cisplatin treatment, which does not depend on mitotic arrest, was hardly affected by the level of Survivin. These data strongly indicate that by compromising the SAC Survivin depletion facilitates mitotic slippage, thereby enhances cell survival against Docetaxel.

Note that cell with a defective SAC does not necessarily develop resistance against taxanes, for other mechanisms unrelated to mitotic arrest may also contribute to treatment outcomes. Inhibition of the SAC kinase Mps1 was reported to enhance the anti-tumor efficacy of docetaxel. The combined treatments lead to severe chromosome mis-segregation rather than mitotic arrest ^44^. Therefore, the role of SAC in cellular response to taxanes remains controversial. Whether depletion of Survivin promote chromosome segregation errors induced by taxanes requires further investigation.

In addition to apoptosis, Survivin inhibition was also reported to cause mitotic catastrophe, which is now recognized as a process preceding apoptosis, necrosis or senescence. Accordingly, we find Survivin-depleted cells showed elevated level of mitotic catastrophe and cellular senescence following Docetaxel treatment. Cellular senescence is an irreversible program of cell-cycle arrest that can be induced by diverse stimuli. Tumors with inactivated apoptotic signaling pathway retain the capacity to become senescent ^45^. Therefore, selectively induction of cellular senescence could represent a promising therapeutic strategy ^46^. Moreover, we found the combination of Survivin depletion and Docetaxel treatment almost completely blocked colony formation of tumor cells. Since clonogenic assay measures the net result of multiple forms of cell death and examines the long-term cellular response that are more reflective of therapeutic response ^35^. Our observations that depletion of Survivin switches Docetaxel induced apoptosis to senescence and enhances the efficacy of Docetaxel against clonogenic survival of tumor cells suggest that non-apoptotic mechanisms may be more relevant to the long-term effect of taxanes.

## Material and Methods

### Plasmid Construction

The coding sequence of Survivin and Bcl-xL were cloned from cDNA of HeLa cells by polymerase chain reaction. DNA encoding Bcl-2 was synthesized and base optimized without changing amino acids owing to a high GC content by Sangon Biotech Co., Ltd. These coding sequences were first cloned into pcDNA3.1 (Invitrogen), whereupon site-directed mutagenesis was performed to generate phosphomimetic or deficient mutants of each gene. The WT or mutant sequences were then cloned into the lentiviral vector pLVX-Puro (Clontech, Mountain View, CA) for lentivirus production. The luciferase gene used for the control was cloned from the pGL3-Basic Vector (Promega, Madison, WI). All inserts were confirmed by DNA sequencing. All primer sequences are listed in Table 1.

**Table 1.**
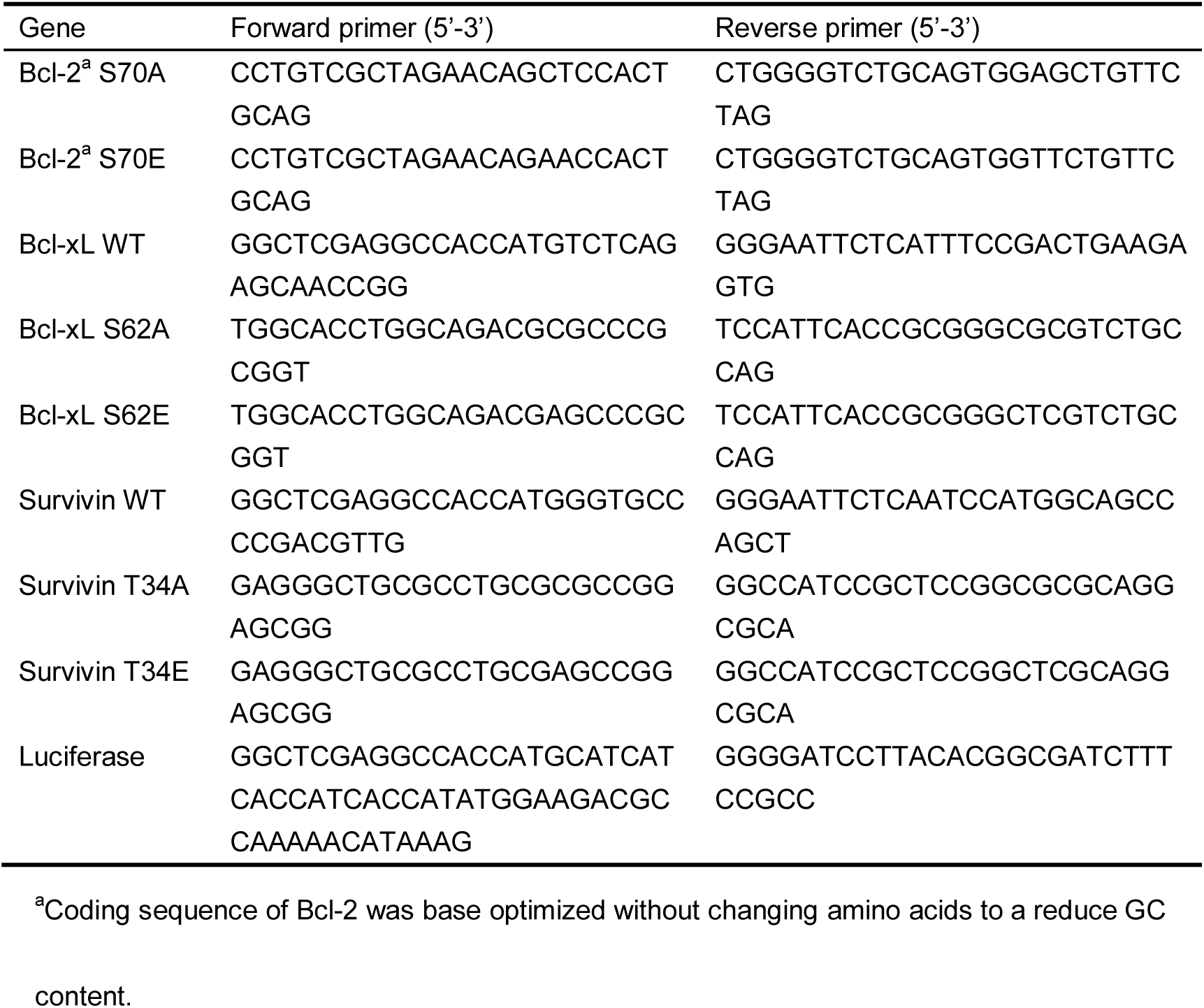
Primer used for vector construction.

### Lentivirus Production

Lentiviral particles were produced according to the “pLKO.1 Protocol” provided by Addgene (Cambridge, MA). Briefly, WT and mutant sequences of Survivin, Bcl-2, and Bcl-xL were individually cloned into the pLVX-Puro vector. The resultant lentiviral vectors were co-transfected with the packaging plasmid psPAX2 (a gift from Didier Trono, Addgene plasmid # 12260) and envelope plasmid pMD2.G (a gift from Didier Trono, Addgene plasmid # 12259) into 293T cells using Lipofectamine 2000 (Invitrogen). After 12–15 h incubation, the medium was replaced with fresh DMEM + 10% FBS. Lentiviral particle-containing medium was harvested from cells after 48 h incubation and filtered through a 0.45-μm filter to remove the 293T cells, then was directly used to infect target cells.

### Cell Culture and Drug Treatment

HeLa, MDA-MB-231, MCF-7, and 293T cells were obtained from the Cell Bank of the Chinese Academy of Sciences (Shanghai, China). HeLa and 293T cells were maintained in Dulbecco’s modified Eagle’s medium (DMEM) (Macgene Technology Ltd., Beijing, China) supplemented with 10% fetal bovine serum (FBS) (Gibco BRL, Life Technologies, Grand Island, NY). MDA-MB-231 cells were grown in L-15 medium (Macgene Technology Ltd.) containing 10% FBS. MCF-7 cells were maintained in DMEM with 10% FBS and 10μg/ml insulin.

Cell lines stably expressing WT and mutant forms of Survivin, Bcl-2, and Bcl-xL were generated by infection with the indicated lentiviral particles, generated as described above. After selection in medium containing μg/ml puromycin for a week, the uninfected cells were no longer viable and overexpression of Survivin, Bcl-2, and Bcl-xL was confirmed by western blot. The expression of mutant protein was confirmed by cDNA pyrosequencing as described below.

For drug treatment, cells were treated with various concentrations of docetaxel (Sanofi-Aventis, Gentilli, France) and Cisplatin (Sigma, St. Louis, MO) and harvested at the indicated time point for analysis.

### Pyrosequencing

Total RNA was isolated from stably transduced cell lines using RNAiso Plus (TaKaRa, Dalian, China) and reverse-transcribed with a PrimeScript™ cDNA Synthesis Kit (TaKaRa) according to the manufacturer’s instructions. Pyrosequencing of each cDNA was performed by Sangon Biotech Co., Ltd. (Shanghai, China).

### Cell Synchronization and Mitotic Index Assessment

Cells were treated with 2 mM thymidine for 20 h followed by incubation in thymidine-free medium for 12 h and a second treatment with 2 mM thymidine for 20 h. Blocked cells were released by washing twice followed by incubation in thymidine-free medium containing 10% FBS.

At the indicated time points after docetaxel treatment, the cells were detached from the cell culture plate using 0.25% trypsin, cytospun, fixed with methanol, and stained with Giemsa solution. The percentage of cells exhibiting condensed chromatin was designated as the mitotic index.

### Small Interfering RNA Transfection

siRNA sequences from previous reports were used to target Survivin (Survivin siRNA #4) ^21^ and Bcl-xL (Bcl-xL siRNA) ^47^. Survivin siRNA #1 was generated using siRNA Wizard^TM^ version 3.1 (InvivoGen, San Diego, CA). siRNA sequences including Survivin siRNA #2, #3 and the negative control siRNA were designed by GenePharma. The 2′-O-methyl modifications of the siRNAs were designed according to previous studies ^31,32^. siRNAs #2m and #3m contained 2’-O-methyl modifications of positions 1 + 2 of the sense strand and position 2 of the guide strand. The siRNA AS4U contained 2’-O-methyl modification of 4 uridine residues in the seed region of the antisense strand. The siRNA AS8U contained 2’-O-methyl modification of all 8 uridine residues in the antisense strand. The siRNA AS1G contained 2’-O-methyl modification of 1 guanosine residue in the antisense strand. Mismatch designs C6-C10 were generated by replacing bases 6–10 with the complement of the original siRNA, respectively. siRNA transfection was performed using the Lipofectamine RNAIMAX reagent (Invitrogen) according to the manufacturer’s instructions. The final siRNA concentration used in all experiments was 40 nM. All siRNA sequences are listed in Table 2.

**Table 2.**
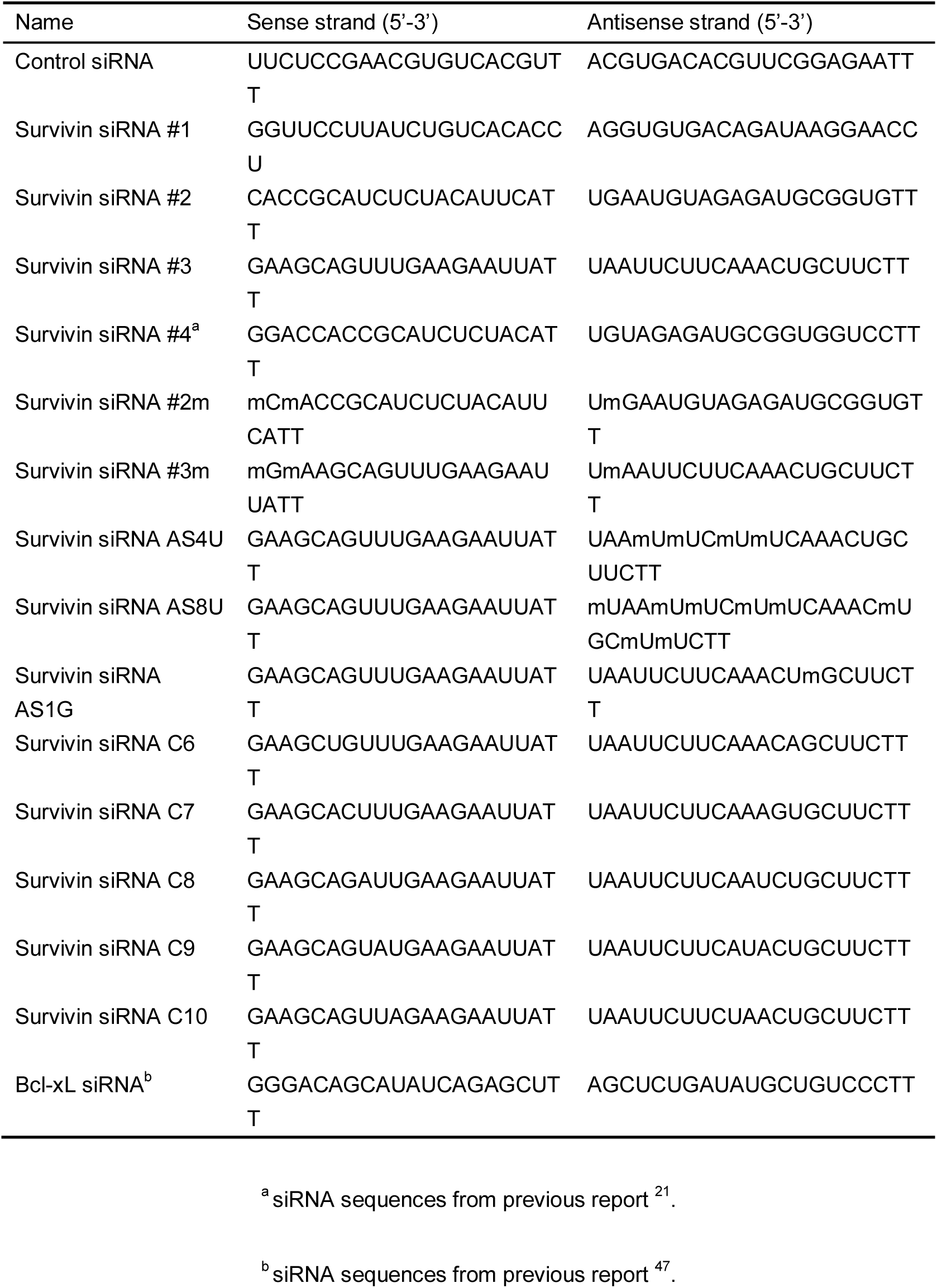
siRNA used for protein depletion.

### Cell Viability Assay

A Cell Counting Kit-8 (CCK-8) (Dojindo, Kumamoto, Japan) was used to assess cell viability according to the manufacturer’s instructions. Cell viability was determined by measuring the absorbance at 450 nm using a microplate reader.

### FACS Apoptosis Assay

Annexin V-FITC/PI staining was performed on fresh cells according to the manufacturer’s specifications (Dojindo). The percentages of apoptotic cells were quantified by combining both early (Annexin V+/PI−) and late (Annexin V+/PI+) apoptotic cells.

### Western blot analysis

Cell pellets were lysed in RIPA buffer (Macgene Technology Ltd., Beijing, China) with 10μl/ml protease inhibitor cocktail (P8340, Sigma) and 10μl/ml phosphatase inhibitor cocktail (p0044, Sigma). The proteins in cell lysates were separated by electrophoresis on a 15% SDS-polyacrylamide gel, transferred to nitrocellulose membranes, immunostained, and visualized by enhanced chemiluminescence detection reagents (Applygen Technologies Inc., Beijing, China). Antibodies against Survivin, Bcl-2, Bcl-xL, P-Histone 3, phospho-Survivin, and phospho-Bcl-2 were purchased from Cell Signaling (Danvers, MA), and the antibody specific for phospho-Ser-62-Bcl-xL was purchased from Abcam (Cambridge, UK). The β-Actin antibody was purchased from Zsbio (Beijing, China).

### Nuclear staining of mitotic cells

Synchronized cells were blocked in mitosis by Docetaxel treatment. Hoechst 33342 (#4082 Cell Signaling) solution was added to the growth medium to obtain a final concentration of 1 μg/ml and incubated at 37□C for 20 min. Cells were then observed and photographed under a fluorescence microscopy (ZEISS Axio Vert.A1).

### SA-β-Gal assay

Cells were seeded in 12-well plates. Following indicated treatments, cells were allowed to grow in drug-free medium for more than 1 week. Senescent cells were detected by using Senescence β-Galactosidase Staining Kit (#9860 Cell Signaling) following the manufacturer’s instructions.

### Colony formation assays

Cells were seeded in 12-well plates. Following treatments, cells were allowed to grow in drug-free medium for 3 weeks. And then fixed with methanol, and stained by crystal violet. The colony is defined to consist of at least 50 cells.

## Acknowledgments

We thank Randy Schekman and Tony Hunter for their guidance.

## Author contributions

T.L.H. designed the research, performed all experiments and wrote the manuscript. H.S. provided expertise and feedback. Y.T.L., D.S.W., J.J., H.L. and B.H. supported the research. Z.X.J. provided funding and provided critical advice.

## Data availability

The datasets used or analysed during the current study are available from the corresponding author on request.

